# DualBranchEEG: A Dual-Branch CNN–BiLSTM Network with Temporal Attention and Spectral Features for Three-Class EEG Seizure Classification

**DOI:** 10.64898/2026.11.25.768266

**Authors:** Ranveer Singh, Jaideep Varma, Aryan Somanna, Shrey Jauhari

## Abstract

Automated seizure detection from electroencephalography (EEG) is a key step toward closed-loop brain–computer interface (BCI) systems and clinical decision support. We present DualBranchEEG, a dual-branch network that fuses (i) a temporal branch—a three-block 1-D convolutional encoder, a two-layer bidirectional LSTM and an additive attention layer— operating on the raw z-scored waveform, with (ii) a spectral branch—an MLP on log-power spectral density (PSD) features from Welch’s method. We evaluate three-class classification (normal, interictal, ictal) on the Bonn University EEG dataset (500 single-channel recordings, 173.61 Hz) using a single stratified 70/15/15 split (seed 42). We position this work as a transparent reference implementation and exploratory comparison, not as evidence of superiority. A CNN–BiLSTM baseline (728,387 parameters; last-time-step readout, no attention, no augmentation) attains 96.00% test accuracy (72/75) and a macro AUC-ROC of 0.9795; five-fold stratified cross-validation of this baseline gives 98.40 *±* 1.52% accuracy and 0.9952 *±* 0.0071 AUC-ROC. The full DualBranchEEG (823,556 parameters) attains the same 96.00% test accuracy, macro F1 of 0.9543 and macro AUC-ROC of 0.9952. In the ablation, CNN-only, BiLSTM-only and CNN+BiLSTM+attention variants each reach 94.67%, and the full model is one test recording better; with only 75 test recordings these differences lie within sampling uncertainty (95% Wilson interval for 96.00%: 88.9–98.6%). Band-power differences between all class pairs are significant in all five canonical EEG bands. Grad-CAM, t-SNE and attention visualisations are reported as qualitative diagnostics only. We document all known methodological limitations, including PSD standardisation fitted on all 500 recordings and Welch’s default 50% segment overlap.

## 1 Introduction

Epilepsy affects approximately 50 million people worldwide, making it one of the most prevalent neurological disorders [1]. Clinical diagnosis and seizure monitoring rely heavily on visual inspection of EEG by expert neurologists—a process that is time-consuming, subjective and impractical for continuous long-term monitoring. Automated seizure detection therefore plays an essential role in modern neurological care and in emerging BCI applications such as closed-loop neurostimulation.

EEG signals are non-stationary, vary across and within patients, are contaminated by muscular and ocular artefacts, and contain both transient events (spikes, high-frequency oscillations) and sustained rhythms (delta, theta, alpha, beta, gamma). An effective detector must capture fine-grained local temporal structure and longer-range rhythmic dynamics while exploiting the frequency-domain separability of different brain states.

Traditional pipelines combine hand-crafted features (wavelet coefficients, band-power ratios, entropies) with classical classifiers [2, 3, 4]. They are interpretable but require expert feature engineering and often generalise poorly across subjects and recording conditions. Deep 1-D CNNs learn local temporal motifs directly from raw signals [5], while recurrent networks model temporal dependencies [6, 7]. Most existing deep models, however, operate on either the raw waveform or a pre-computed spectral representation, leaving the complementarity of the two views largely unexploited.

### 1.1 Contributions

- **Dual-branch architecture.** DualBranchEEG jointly encodes the raw waveform (CNN*→*BiLSTM*→*additive attention) and Welch log-PSD features (MLP), fused by concatenation before a shared classi-fier head.
- **Temporal attention.** A lightweight additive attention layer replaces last-time-step readout of the BiLSTM and yields a per-time-step weight that can be inspected.
- **Controlled comparison with disclosed limitations.** We report a baseline, three ablation variants and the full model on one fixed split, together with exact parameter counts, confidence intervals and a list of every deviation between the intended and the executed protocol (Sections 7 and 11).
- **Qualitative diagnostics.** Grad-CAM, t-SNE and attention overlays, and a Mann–Whitney band-power analysis, are provided as descriptive evidence; none is presented as clinical validation.

## 2 Related Work

### 2.1 Classical Feature-Based Methods

Early automated detectors relied on rule-based analysis of EEG morphology [2] and on features extracted from wavelet sub-bands, fed to neural-network or logistic-regression classifiers [3]. Non-linear measures such as approximate, sample and phase entropy provide complementary discriminative information [4]. These methods remain useful interpretable baselines but require careful feature engineering.

### 2.2 Convolutional Neural Networks

Acharya et al. [5] applied a 13-layer 1-D CNN to raw Bonn EEG for normal, pre-ictal and seizure classes and reported 88.67% accuracy under ten-fold cross-validation, eliminating manual feature extraction. Ullah et al. [8] later proposed an ensemble of pyramidal 1-D CNNs for Bonn classification, and Wei et al. [11] applied 3-D CNNs to multi-channel EEG for seizure detection. Class definitions, data subsets and validation schemes differ across such studies, so reported accuracies are not directly comparable.

### 2.3 Recurrent and Attention-Based Models

LSTM networks have been applied to seizure prediction from EEG [6] and to seizure detection [7]. Bidirectional recurrence [9] lets each hidden state use both past and future context, which is attractive for offline classification of fixed-length segments. Transformer-based self-supervised models such as BENDR [10] replace recurrence by self-attention, at quadratic cost in sequence length.

### 2.4 Hybrid CNN–RNN Architectures

Combining convolutional feature extraction with recurrent modelling has become a common paradigm for biosignal classification. DualBranchEEG extends the CNN–BiLSTM design with an attention readout and a complementary spectral branch, evaluated on the three-class Bonn problem.

## 3 Dataset Audit and Preprocessing

### 3.1 Bonn University EEG Dataset

We use the public dataset of Andrzejak et al. [12]: five sets (Z, O, N, F, S) of 100 single-channel segments, each 23.6 s long (4097 samples) at 173.61 Hz with 12-bit resolution. The three-class mapping is shown in Table 1. Loading in the experiment notebook confirmed 100 files per set, a data array of shape (500, 4097) and class counts of 200 / 200 / 100 (normal / interictal / ictal).

**Table 1.**
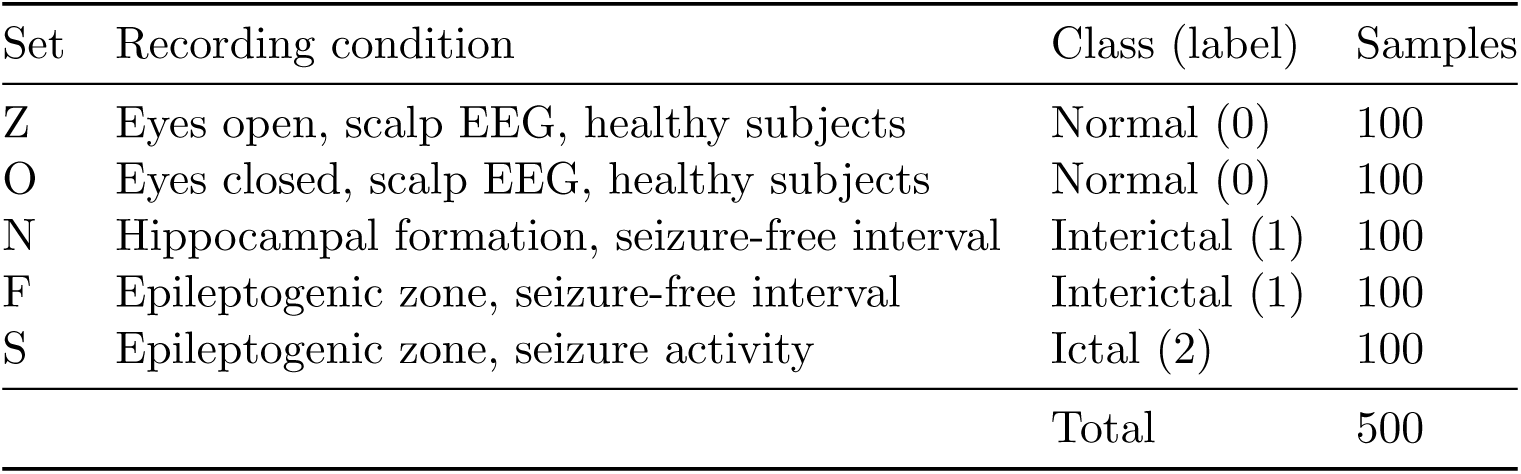
Bonn University EEG dataset audit. Class mapping used throughout.

| Set | Recording condition | Class (label) | Samples |
|---|---|---|---|
| Z | Eyes open, scalp EEG, healthy subjects | Normal (0) | 100 |
| O | Eyes closed, scalp EEG, healthy subjects | Normal (0) | 100 |
| N | Hippocampal formation, seizure-free interval | Interictal (1) | 100 |
| F | Epileptogenic zone, seizure-free interval | Interictal (1) | 100 |
| S | Epileptogenic zone, seizure activity | Ictal (2) | 100 |
|  |  | Total | 500 |

The resulting class distribution (Normal 200, Interictal 200, Ictal 100) is mildly imbalanced and is handled with a class-weighted loss (Section 4.4).

### 3.2 Exploratory Analysis

Table 2 reports the mean integrated band power per class over the five canonical bands (trapezoidal integration of the Welch PSD, computed on the raw, un-normalised signals). Ictal recordings show markedly higher power in delta, theta, alpha and beta. Interictal recordings have more delta and theta power than normal recordings but less alpha, beta and gamma. Two caveats apply: the Bonn recordings were band-pass filtered (0.53–40 Hz) by the original authors [12], so gamma-band power (30–60 Hz) largely reflects filter roll-off and residual noise; and the Normal class was recorded with scalp electrodes while the Interictal and Ictal classes are intracranial, a confound that may inflate class separability independently of pathology.

### 3.3 Normalisation and Data Splitting

Each segment is standardised independently by per-sample z-scoring,

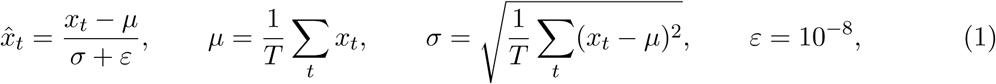

which gives amplitude invariance and uses no information from other recordings. The 500 recordings are split with stratification (seed 42) into training (70%, *n* = 350), validation (15%, *n* = 75) and test (15%, *n* = 75) sets. The baseline and the dual-branch experiments use the same seed and stratification and therefore the same partitions.

**Figure 1.**
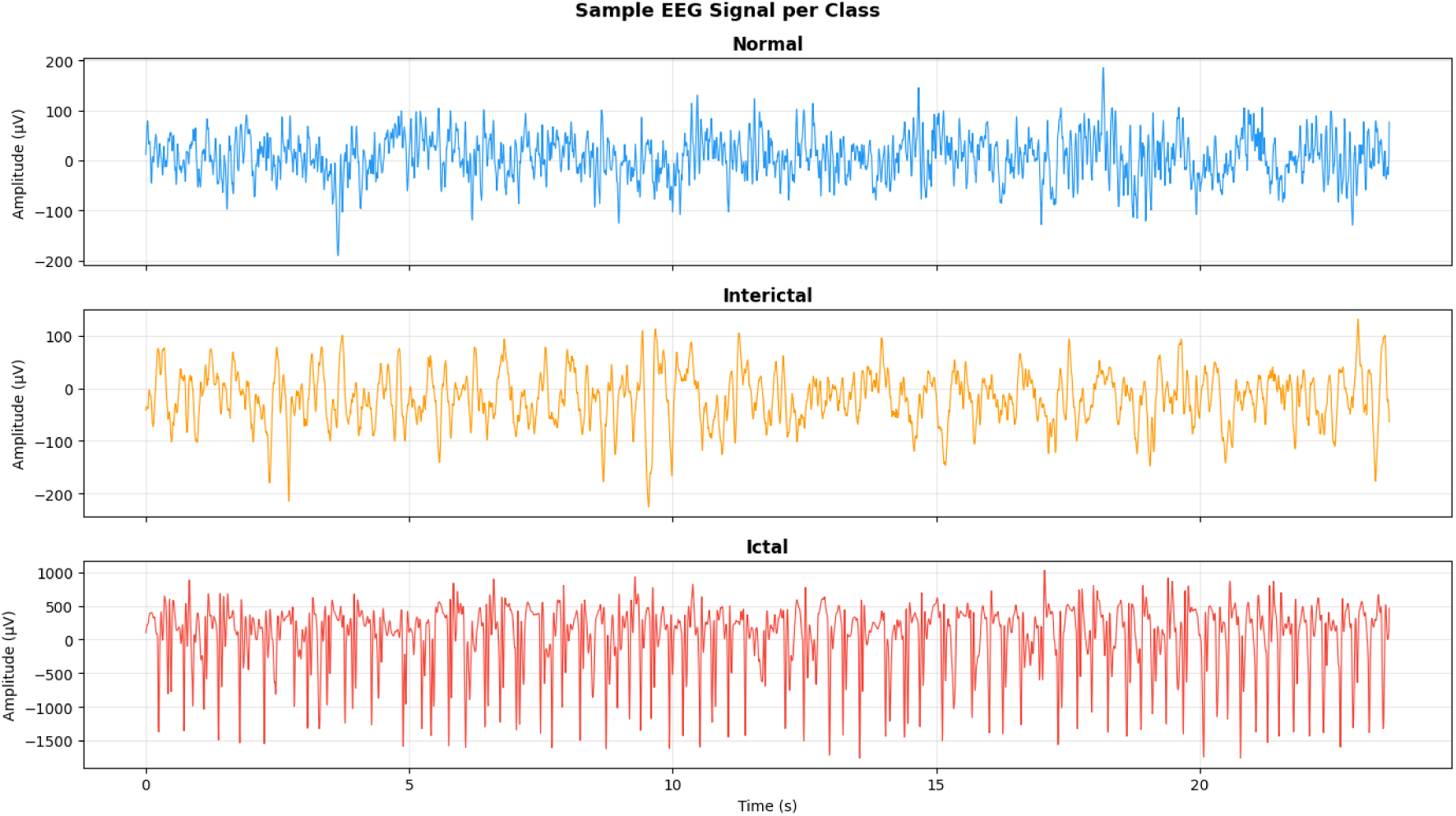
Representative EEG segments. First recording of each class (Normal, Interictal, Ictal), raw amplitude in *µ*V.

**Figure 2.**
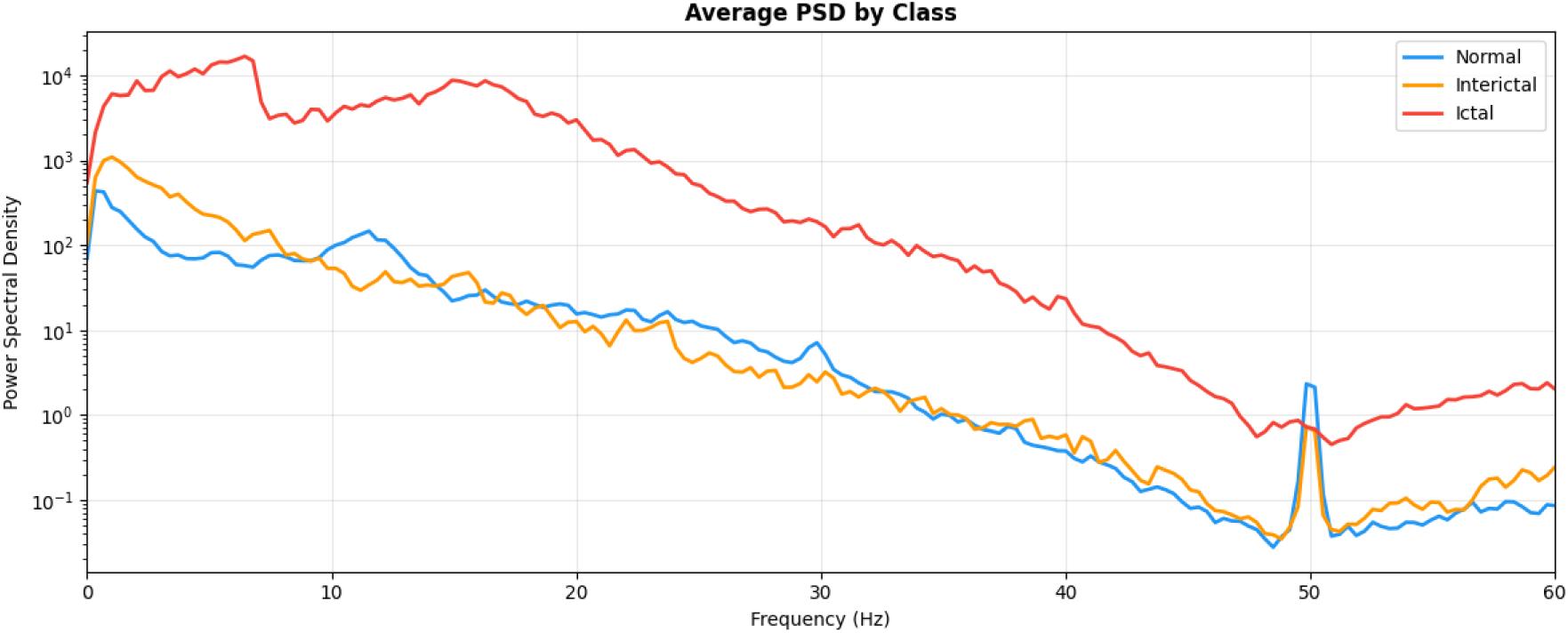
Average Welch PSD by class. Mean over the first 10 recordings of each class (log scale, 0–60 Hz).

**Table 2.** Mean integrated band power (*µ*V^2^) per class (all 500 recordings).

| Class | *δ* (0.5–4) | *θ* (4–8) | *α* (8–13) | *β* (13–30) | *γ* (30–60) |
|---|---|---|---|---|---|
| Normal | 543.9 | 321.5 | 1198.2 | 478.0 | 25.3 |
| Interictal | 2929.6 | 811.9 | 333.9 | 178.1 | 11.3 |
| Ictal | 21273.5 | 41970.9 | 19561.1 | 24939.0 | 438.9 |

### 3.4 Spectral Feature Extraction

PSD estimates are obtained with Welch’s method [13] (SciPy welch, nperseg=512, sampling rate 173.61 Hz). The number of overlapping samples was not set explicitly, so the SciPy default of 50% overlap (256 samples) was used. Bins up to 60 Hz are retained, giving *D*_psd_ = 177 features, which are transformed by log(1 + PSD) and then standardised per frequency bin. This per-bin standardisation was **fitted on all 500 rec ordings before the train/validation/test split and therefore uses unlabeled test-set spectra** (Section 11).

### 3.5 Data Augmentation

During training of the dual-branch model and of the ablation variants (not the baseline), one or two of the following augmentations are drawn at random for each sample:

- Gaussian noise with standard deviation *rσ_x_*, *r ∼ U* (0.05, 0.15) (SNR approximately 16.5– 26 dB).
- Circular temporal shift of up to *±*10% of the segment length.
- Amplitude scaling by *s ∼ U* (0.8, 1.2).
- Crop-and-pad: retain 85–99% of the signal from a random onset and pad the remainder with edge values.

The segment is re-standardised after augmentation. In the dual-branch model augmentation is applied to the raw waveform only; the precomputed PSD vector is not recomputed from the augmented signal.

## 4 Method

Figure 3 gives an overview. The model takes the standardised waveform **x** *∈* R*^T^ ^×^*^1^ (*T* = 4097) and the log-PSD vector **p** *∈* R^177^.

### 4.1 Temporal Branch

**1-D CNN encoder.** Three blocks, each Conv1D, batch normalisation [14], GELU [15], max- pooling (stride 4) and dropout (0.2), with configuration in Table 3. The temporal length is reduced from 4097 to 64 (4097 *→* 1024 *→* 256 *→* 64) and the channel dimension grows to 128, giving **F** *∈* R^64^*^×^*^128^.

**Figure 3.**
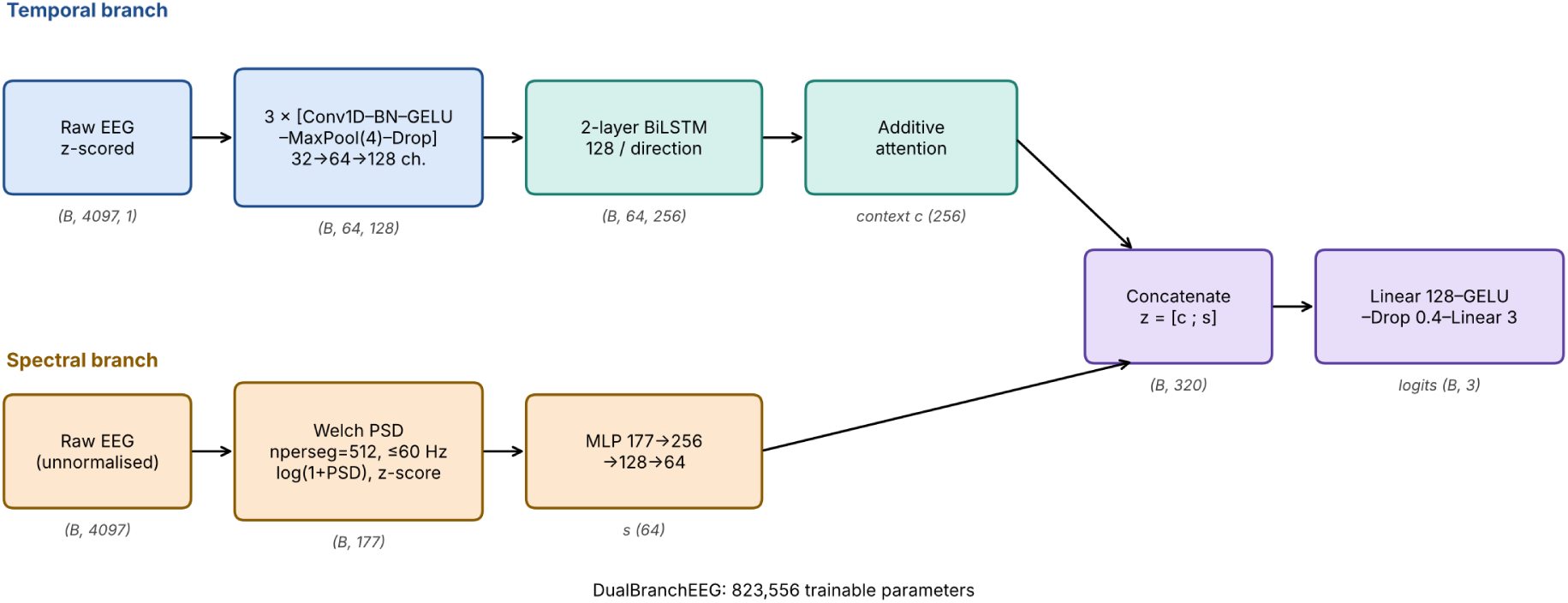
DualBranchEEG architecture. The temporal branch (top; blue = CNN encoder, teal = recurrent/attention stage) encodes the raw waveform; the spectral branch (bottom; orange) encodes log-PSD features. The 256-d context vector and 64-d spectral embedding are concatenated (purple) and classified. Tensor shapes are given for batch size *B*.

**Table 3.**
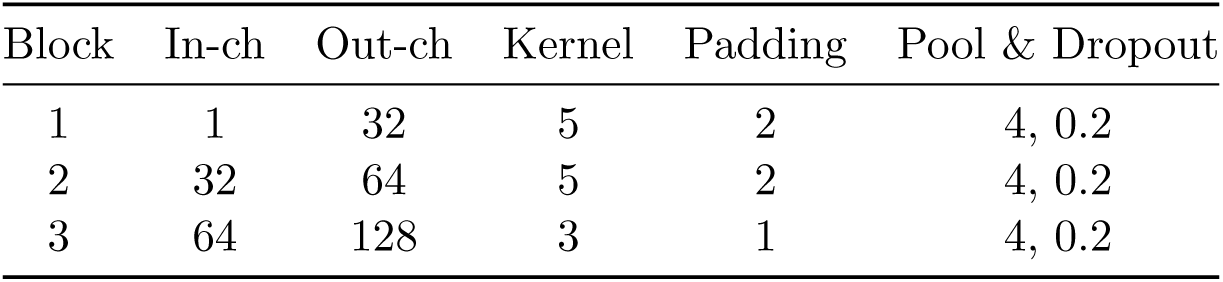
1-D CNN encoder configuration.

| Block | In-ch | Out-ch | Kernel | Padding | Pool & Dropout |
|---|---|---|---|---|---|
| 1 | 1 | 32 | 5 | 2 | 4, 0.2 |
| 2 | 32 | 64 | 5 | 2 | 4, 0.2 |
| 3 | 64 | 128 | 3 | 1 | 4, 0.2 |

**Bidirectional LSTM. F** is passed to a two-layer BiLSTM [9] with hidden size 128 per direction (256 in total) and inter-layer dropout 0.3:

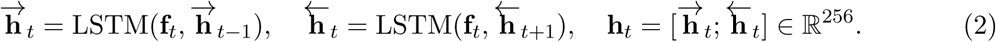

**Additive attention.** The sequence is aggregated by Bahdanau-style additive attention [16]:

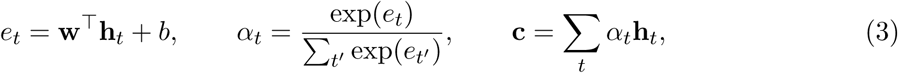

with **w** *∈*R^256^, scalar bias *b* and **c** *∈* R^256^.

**Baseline readout.** The CNN–BiLSTM baseline has no attention layer: it uses the BiLSTM output at the final time step, **h**_64_. In this position the backward direction has processed only the last input step, so the baseline readout under-uses the backward pass; this motivates the attention readout.

### 4.2 Spectral Branch

The log-PSD vector is processed by a three-layer MLP with GELU activation after every linear layer (including the last) and dropout 0.3 after the first two:

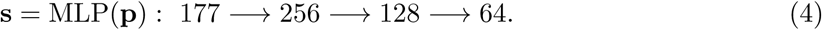

### 4.3 Fusion and Classification

**z** = [**c**; **s**] *∈*R^320^ and **y**^ = *W*_2_ GELU(*W*_1_**z** + **b**_1_) + **b**_2_ with *W*_1_ *∈* R^128*×*320^, *W*_2_ *∈* R^3*×*128^ and dropout 0.4 between the layers.

### 4.4 Loss Function

Class-weighted cross-entropy, with weights computed from the training split:

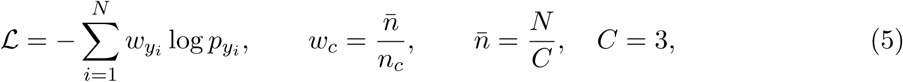

giving *w*_0_ = *w*_1_ = 0.833 and *w*_2_ = 1.667 for Normal, Interictal and Ictal.

## 5 Model Complexity

Parameter counts were derived from the module definitions and agree with the counts printed by the code (728,387 for the baseline; 823,556 for DualBranchEEG).

**Table 4.** Trainable parameters per module. The BiLSTM dominates the budget.

| Module | Parameters | Share of DualBranchEEG |
|---|---|---|
| CNN encoder (3 blocks, incl. BN) | 35,648 | 4.3% |
| BiLSTM (2 layers, bidirectional) | 659,456 | 80.1% |
| Additive attention | 257 | <0.1% |
| Spectral MLP | 86,720 | 10.5% |
| Classifier head | 41,475 | 5.0% |
| DualBranchEEG total | 823,556 | 100% |
| CNN–BiLSTM baseline (no attention, no spectral branch) | 728,387 | – |

The approximately 0.82 M-parameter model is small enough for resource-constrained hardware in principle, but inference latency and memory were not profiled in this work.

## 6 Algorithm

### Algorithm 1: DualBranchEEG forward pass

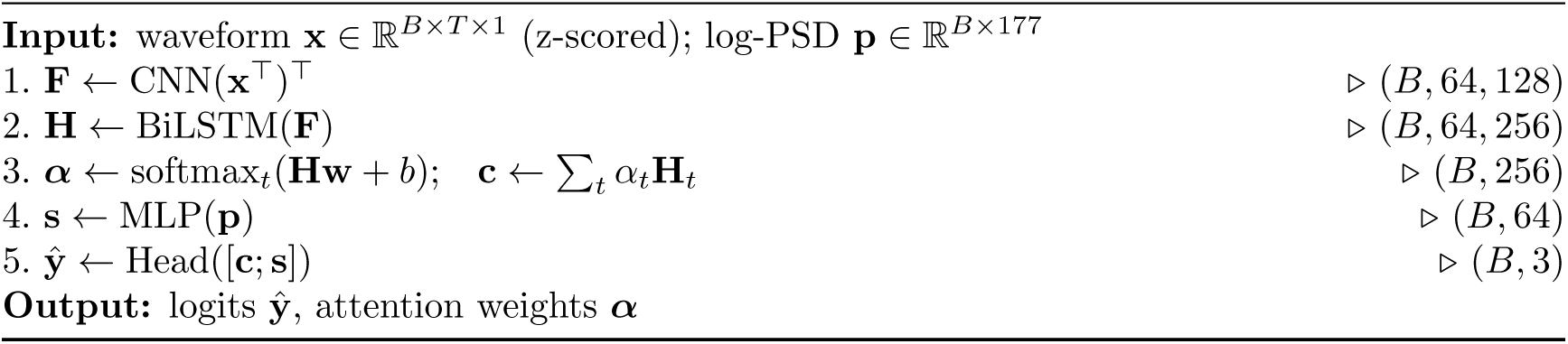

## 7 Experimental Setup

### 7.1 Data Loading and Preprocessing

Recordings are read from the tab-separated text files of each set, converted to float32, z-scored per segment and split as in Section 3.3. PSD features are computed as in Section 3.4.

### 7.2 Training Protocol

All models use AdamW [17] (*β*_1_ = 0.9, *β*_2_ = 0.999, weight decay 10*^−^*^4^), initial learning rate 10*^−^*^3^ with cosine annealing [18] to *η*_min_ = 10*^−^*^5^, gradient-norm clipping at 1.0 and mini-batches of 16 (drop last during training). The checkpoint with the highest validation accuracy is used for testing. Table 5 lists the training budgets, which differ between models.

**Table 5.**
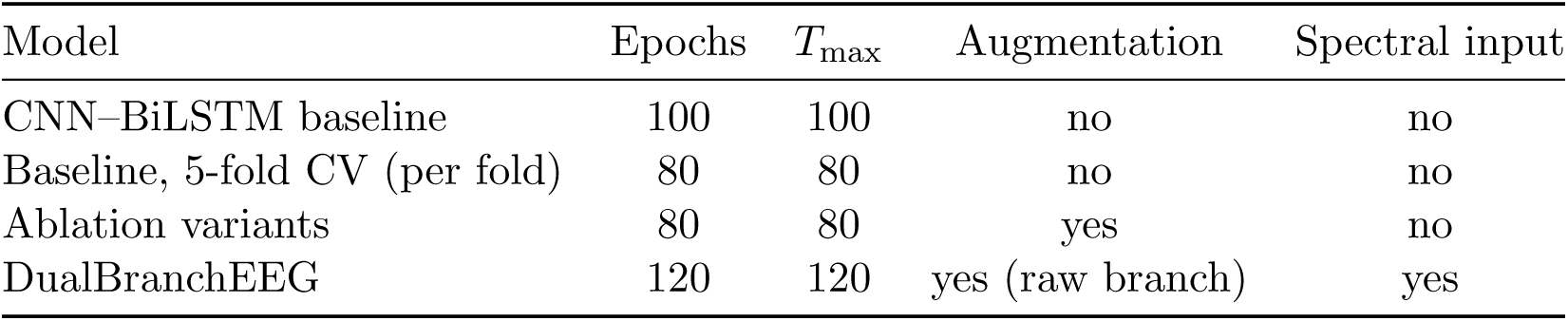
Training configurations as executed.

| Model | Epochs | *T*<sub>max</sub> | Augmentation | Spectral input |
|---|---|---|---|---|
| CNN–BiLSTM baseline | 100 | 100 | no | no |
| Baseline, 5-fold CV (per fold) | 80 | 80 | no | no |
| Ablation variants | 80 | 80 | yes | no |
| DualBranchEEG | 120 | 120 | yes (raw branch) | yes |

A single global seed (42) was set at the start of the notebook and the models were trained sequentially; no repeated-seed runs were performed. Experiments were run on a CUDA GPU in Google Colab; wall-clock times were not recorded.

### 7.3 Evaluation Protocol

Metrics on the held-out test set (75 recordings): accuracy, per-class precision, recall (sensitivity), specificity and F1, macro F1, and one-vs-rest AUC-ROC (macro-averaged). For the baseline we additionally report five-fold stratified cross-validation on all 500 recordings. Confidence intervals are Wilson score intervals [19]. Calibration is summarised by the expected calibration error (ECE) [20] computed one-vs-rest with 10 bins.

### 7.4 Models Compared

i. CNN–BiLSTM baseline, last-step readout; (ii) CNN-only (three blocks, global average pooling, two-layer classifier); (iii) BiLSTM-only (two-layer BiLSTM on the raw waveform, attention, two-layer classifier); (iv) CNN+BiLSTM+Attn (temporal branch of DualBranchEEG without the spectral branch); (v) DualBranchEEG.

## 8 Results

### 8.1 Baseline CNN–BiLSTM

On the held-out test set the baseline reaches 96.00% accuracy (72/75; 95% Wilson interval 88.9–98.6%) and a macro AUC-ROC of 0.9795 (one-vs-rest AUC: Normal 0.977, Interictal 0.979, Ictal 0.982). Three recordings are misclassified. Ictal recordings are classified with perfect precision but the lowest recall (0.87).

**Table 6.** Per-class test performance, baseline CNN–BiLSTM (*n* = 75).

| Class | Precision | Recall | F1 | Support |
|---|---|---|---|---|
| Normal | 0.94 | 1.00 | 0.97 | 30 |
| Interictal | 0.97 | 0.97 | 0.97 | 30 |
| Ictal | 1.00 | 0.87 | 0.93 | 15 |
| Macro avg | 0.97 | 0.94 | 0.95 | 75 |

### 8.2 DualBranchEEG

DualBranchEEG also reaches 96.00% test accuracy (72/75) with macro F1 0.9543 and macro AUC-ROC 0.9952. Accuracy and macro F1 are identical to the baseline; only the AUC-ROC differs (0.9795 vs. 0.9952). Again three recordings are misclassified: one Interictal as Normal and two Ictal as Interictal.

**Table 7.** Per-class test performance, DualBranchEEG (*n* = 75).

| Class | Precision | Recall | Specificity | F1 | AUC | Support |
|---|---|---|---|---|---|---|
| Normal | 0.9677 | 1.0000 | 0.9778 | 0.9836 | 1.0000 | 30 |
| Interictal | 0.9355 | 0.9667 | 0.9556 | 0.9508 | 0.9978 | 30 |
| Ictal | 1.0000 | 0.8667 | 1.0000 | 0.9286 | 0.9878 | 15 |
| Macro avg | 0.9677 | 0.9444 | – | 0.9543 | 0.9952 | 75 |

### 8.3 Five-Fold Cross-Validation (Baseline)

Table 8 gives per-fold results for the baseline. The mean is 98.40*±*1.52% accuracy, 0.9952*±*0.0071 AUC-ROC and 0.9850 *±* 0.0140 macro F1 (sample standard deviation over folds). Two points qualify these numbers: the best epoch of each fold was selected by accuracy on the same fold that is then reported, which biases the estimate upward, and cross-validation was not run for DualBranchEEG. The higher CV accuracy relative to the 96.00% held-out test accuracy is consistent with this bias and with the small test set.

**Table 8.**
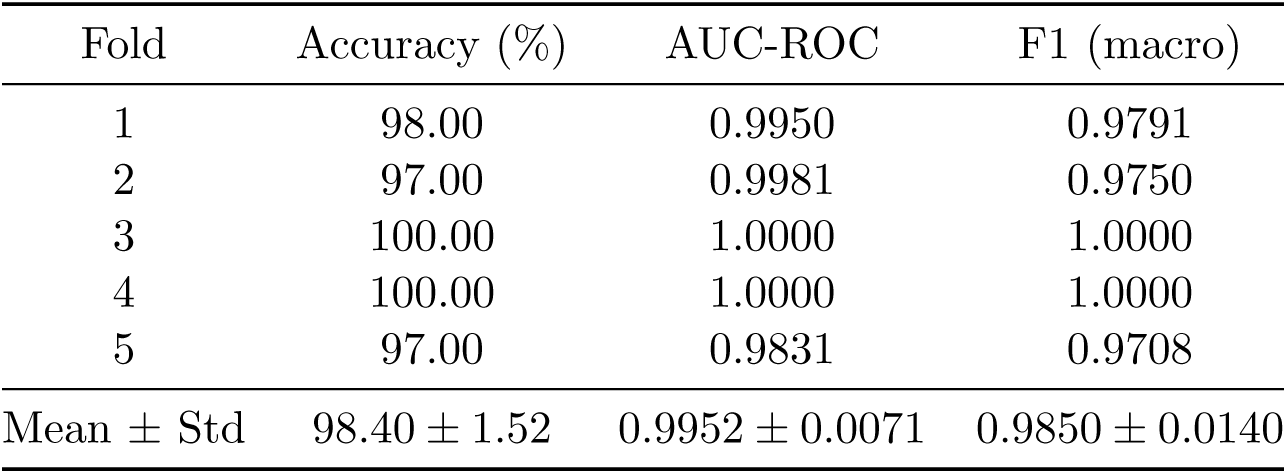
Five-fold stratified cross-validation, baseline CNN–BiLSTM (80 epochs per fold, 100 recordings per fold).

### 8.4 Comparison with Prior Work

We do not tabulate literature accuracies against ours: published Bonn results use different class definitions, subsets, and validation schemes (hold-out, *k*-fold, subject- or segment-level splits), so numerical comparison would be misleading. For orientation, Acharya et al. [5] report 88.67% for normal / pre-ictal / seizure under ten-fold cross-validation, and ensemble CNNs [8] report accuracies near 99% under their own protocol. Our hold-out result (96.00%) and CV result (98.40%) should be read in the context of the limitations in Section 11, and we make no claim of state-of-the-art performance.

## 9 Ablation Study

Table 9 lists test results for four variants and the baseline. The three temporal-only variants tie at 94.67% (71/75); their AUC-ROC and F1 differ slightly. The full model is one recording better in accuracy (96.00%, 72/75) and highest in F1 and AUC-ROC.

**Table 9.**
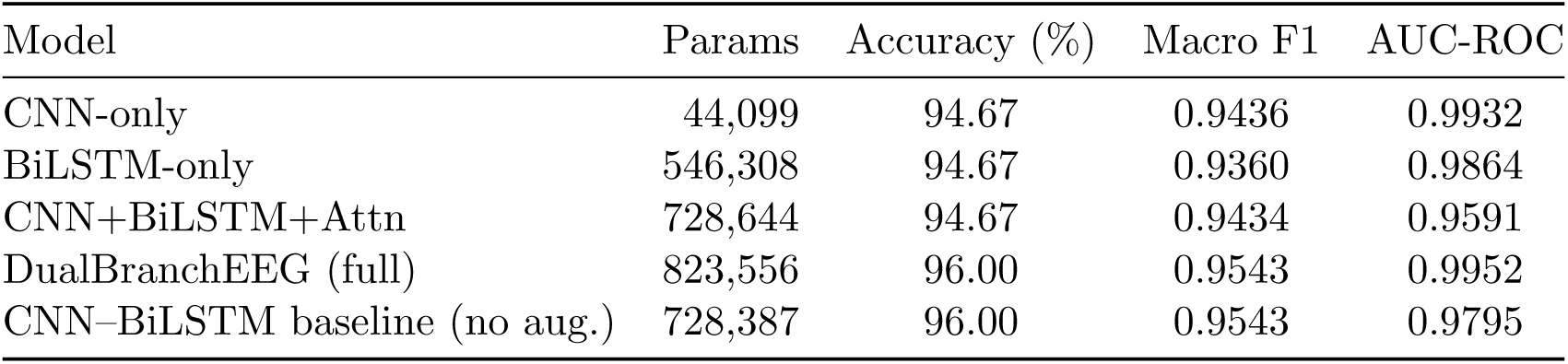
Ablation on the held-out test set (*n* = 75, single seed). Budgets differ (Table 5).

| Model | Params | Accuracy (%) | Macro F1 | AUC-ROC |
|---|---|---|---|---|
| CNN-only | 44,099 | 94.67 | 0.9436 | 0.9932 |
| BiLSTM-only | 546,308 | 94.67 | 0.9360 | 0.9864 |
| CNN+BiLSTM+Attn | 728,644 | 94.67 | 0.9434 | 0.9591 |
| DualBranchEEG (full) | 823,556 | 96.00 | 0.9543 | 0.9952 |
| CNN–BiLSTM baseline (no aug.) | 728,387 | 96.00 | 0.9543 | 0.9795 |

**Figure 4.**
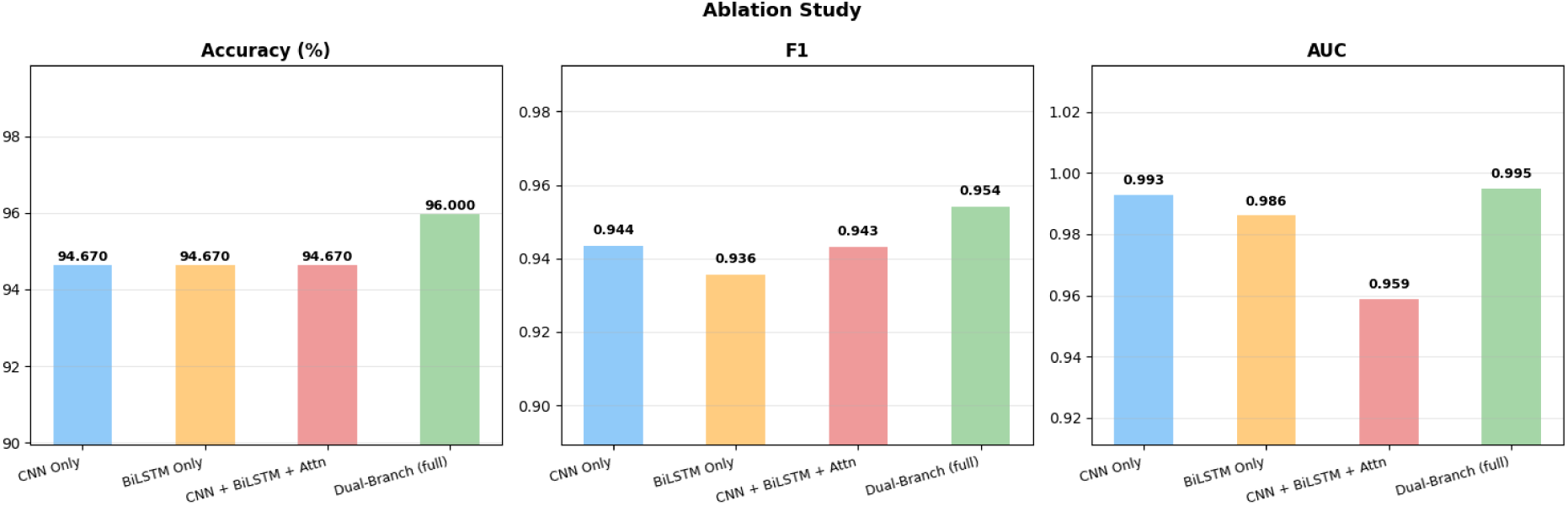
Ablation results. Accuracy, F1 and AUC-ROC for CNN-only, BiLSTM-only, CNN+BiLSTM+Attn and DualBranchEEG.

### Interpretation

The data do *not* support a claim that each component improves performance incrementally. Adding the BiLSTM or attention to the CNN does not change accuracy, and CNN+BiLSTM+Attn has the lowest AUC-ROC (0.9591). BiLSTM-only matches CNN-only in accuracy, so the data do not show that convolutional features are essential. The only improvement is for the full model, and it amounts to one test recording (1.33 percentage points), well inside the 95% interval of either result (94.67%: 87.1–97.9%; 96.00%: 88.9–98.6%). The unaugmented baseline matches the full model’s accuracy, and the full model was trained for more epochs (120 vs. 80), so a contribution of the spectral branch cannot be separated from training budget with this design. A seed-repeated, budget-matched study with paired tests is needed.

## 10 Analysis

### 10.1 Training Dynamics

The baseline reaches 100% training accuracy by epoch 60 while validation accuracy ranges 94.7–98.7% after epoch 20, with a best validation accuracy of 98.67% (reached no later than epoch 50). DualBranchEEG fits the training set almost perfectly by epoch 20 (loss *≈* 3 *×* 10*^−^*^4^); validation loss then rises from 0.046 (epoch 20) to about 0.21 (epoch 120) while validation accuracy plateaus at 94.7–97.3% (best 98.67%), indicating overfitting that checkpoint selection on 75 validation recordings only partly mitigates.

### 10.2 Statistical Analysis of Band Power

Mann–Whitney *U* tests [21] were applied to each of the five bands for each of the three class pairs (15 tests). All 15 comparisons give *p <* 10*^−^*^5^ (uncorrected; all remain significant under Bonferroni correction, threshold 0.05*/*15 *≈* 0.0033). Class-wise spectral differences therefore exist in every band, as visible in Table 2 and Figure 12. This establishes that the PSD features carry class information; it does not by itself show which features the trained network uses.

**Figure 5.**
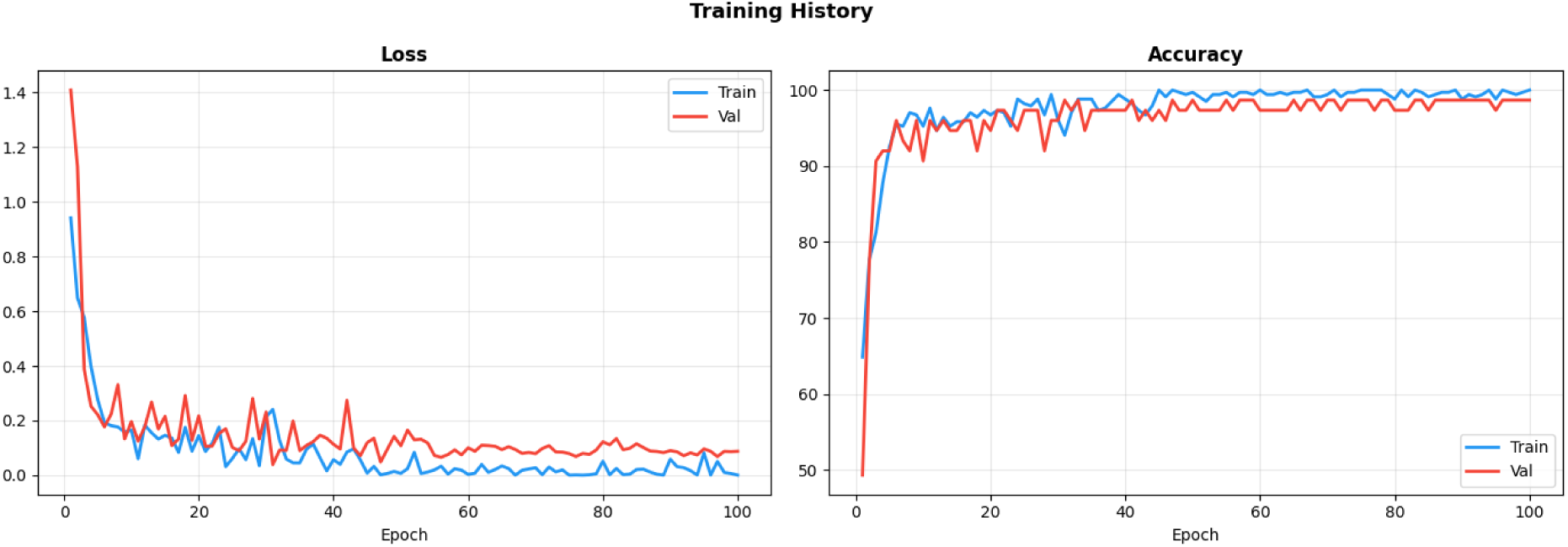
Baseline training history (100 epochs).

**Figure 6.**
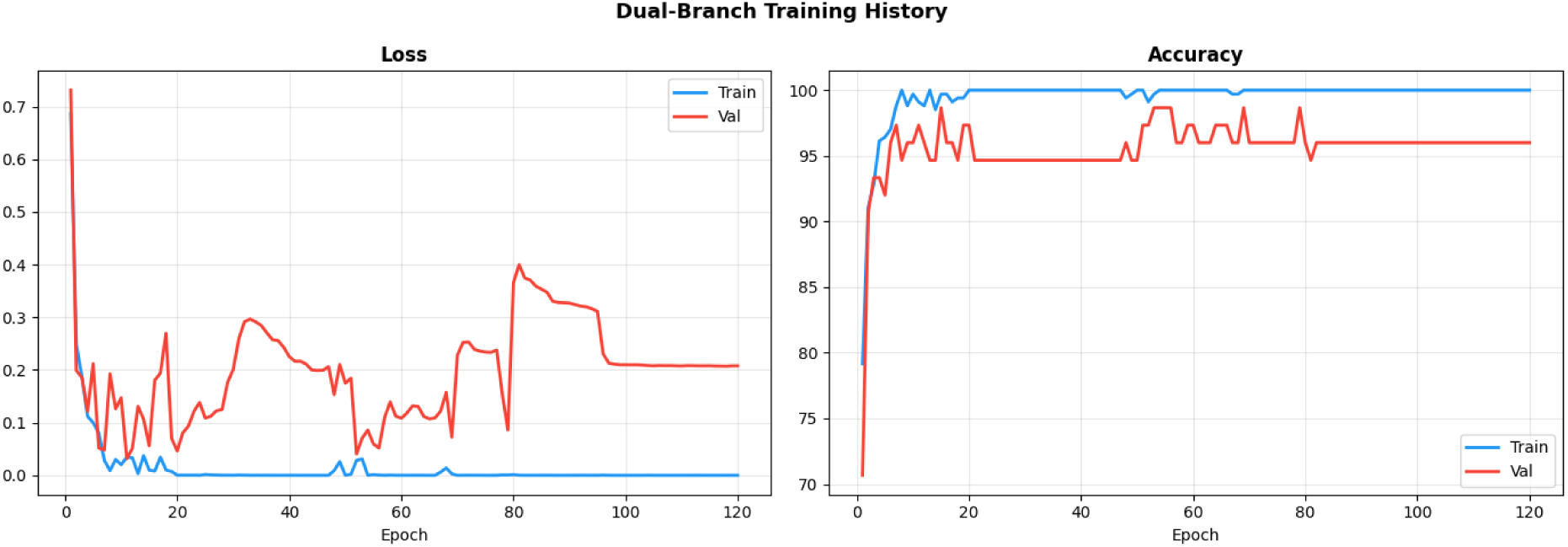
DualBranchEEG training history (120 epochs).

**Table 10.** Mann–Whitney *U* statistics (all *p <* 10*^−^*^5^).

| Band | Normal vs Interictal | Normal vs Ictal | Interictal vs Ictal |
|---|---|---|---|
| *δ* | 8083 | 246 | 2707 |
| *θ* | 12789 | 9 | 242 |
| *α* | 32504 | 956 | 142 |
| *β* | 34418 | 387 | 122 |
| *γ* | 32074 | 1458 | 569 |

### 10.3 Interpretability Diagnostics

#### Grad-CAM

Grad-CAM [22] was computed from the baseline CNN–BiLSTM (not Dual-BranchEEG) at the output of the third convolution (before batch normalisation and activation):

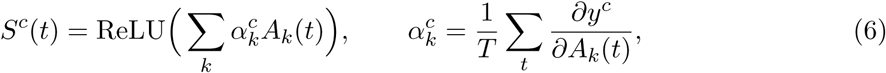

linearly upsampled to 4097 samples and max-normalised. It was applied to the first recording of each class in the full dataset (not a test-only selection), with the network in training mode (dropout active). Saliency is sparse and localised to short bursts for the Normal and Interictal examples and more widely distributed across the high-amplitude rhythmic activity of the Ictal example. We do not claim alignment with specific clinical waveforms from these single examples.

#### t-SNE

The 128-d CNN features (temporal mean of the encoder output of the baseline) of all 500 recordings were projected with t-SNE [23] (perplexity 30, 1000 iterations). The three classes form clearly separated clusters with a few stray Ictal points near the Interictal and Normal clusters. Because the 500 recordings include the 350 training samples seen by the network, the plot shows learned structure rather than generalisation, and t-SNE cannot establish linear separability.

#### Attention weights

Attention weights of DualBranchEEG were plotted for the first test recording of each class (Figure 9). The weights are close to uniform over the 64 time steps (about 0.012–0.017, versus 1*/*64 *≈* 0.0156 for uniform attention), with only slow modulation; they do not concentrate on specific events. The displayed Ictal example is misclassified as Interictal. In this model the attention layer therefore acts almost like average pooling, and we draw no clinical conclusion from the attention maps.

### 10.4 Calibration and Error Analysis

For the baseline, one-vs-rest ECE (10 bins) is 0.0278 (Normal), 0.0322 (Interictal) and 0.0300 (Ictal); with 75 samples and 10 bins these values are noisy. The calibration curves in Figure 13 are plotted with 8 bins. The errors of both models are concentrated at class boundaries: Ictal recordings predicted as Interictal (and, for the baseline, one as Normal), and Interictal recordings predicted as Normal.

## 11 Limitations and Failure Modes

- **Transductive PSD standardisation.** Per-bin standardisation of the PSD features used mean and standard deviation over all 500 recordings, including validation and test samples, before splitting. It uses no labels, but it is a leak of test-set statistics; the effect on reported results has not been measured. Fitting on the training set only and re-running is required.
- **Welch overlap.** The overlap was left at the SciPy default (50%), not the 0% originally intended; reported features correspond to the executed setting.
- **Small, single-split evaluation.** The test set has 75 recordings, each error moves accuracy by 1.33 points, and one seed was run. Differences between models are not statistically established.
- **Unequal training protocols.** The baseline was trained without augmentation for 100 epochs, the ablation variants for 80 and DualBranchEEG for 120 epochs. In DualBranchEEG the spectral branch sees unaugmented PSD vectors.
- **Cross-validation bias.** CV was run for the baseline only, with checkpoint selection on the evaluated fold.
- **Dataset confounds.** Single-channel, pre-filtered (0.53–40 Hz), small and clean; the Normal class is scalp EEG and the other classes are intracranial; segments from the same subject may appear in different splits (subject identifiers are not provided).
- **Interpretability.** Grad-CAM and t-SNE use the baseline model and include training samples or non-test examples; attention is near uniform.
- **Not evaluated.** Multi-channel EEG, large clinical corpora (CHB-MIT [24], TUSZ [25]) and real-time latency.

**Figure 7.**
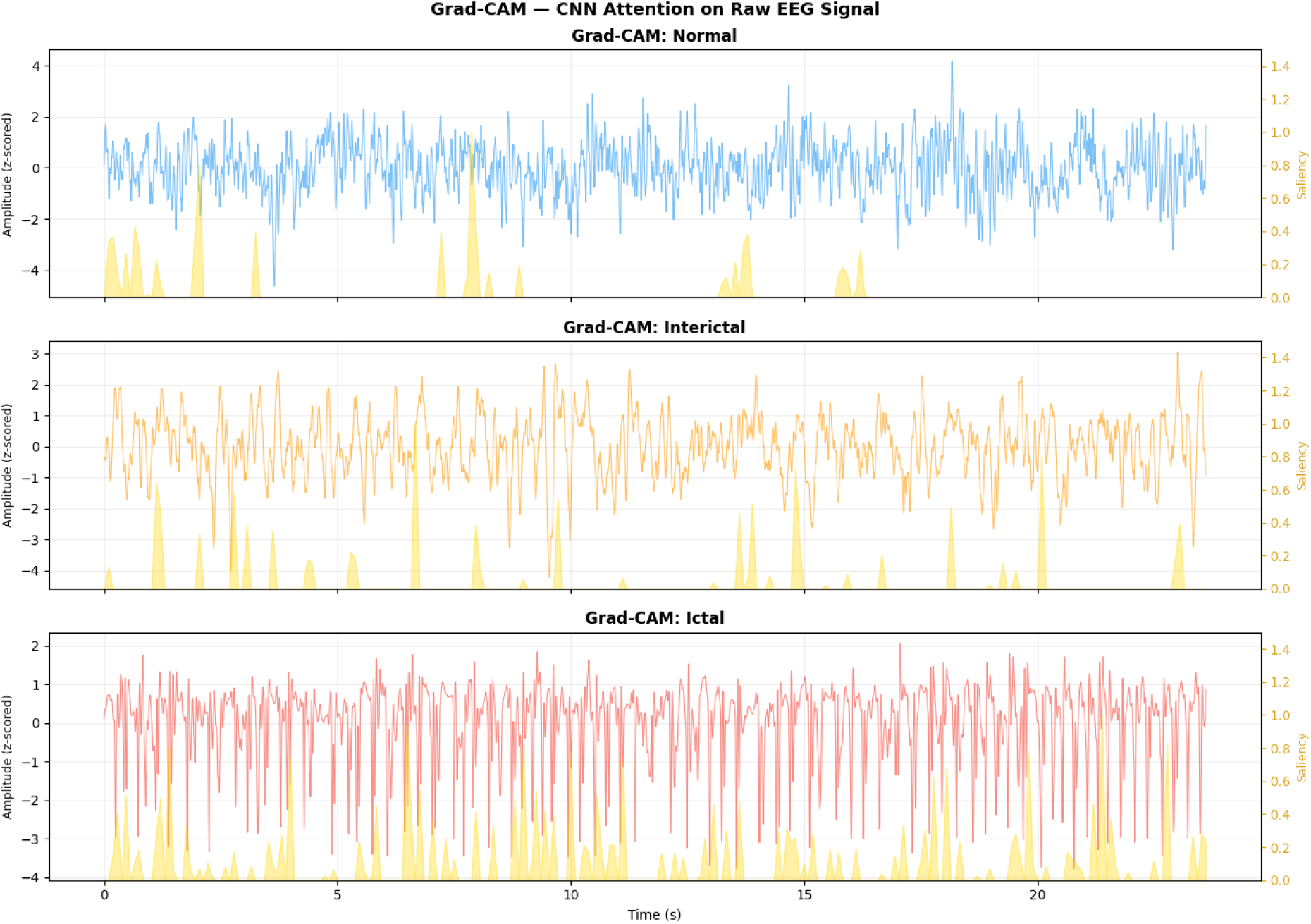
Grad-CAM saliency (gold) overlaid on one z-scored recording per class, baseline CNN– BiLSTM. Single examples; not a quantitative evaluation.

**Figure 8.**
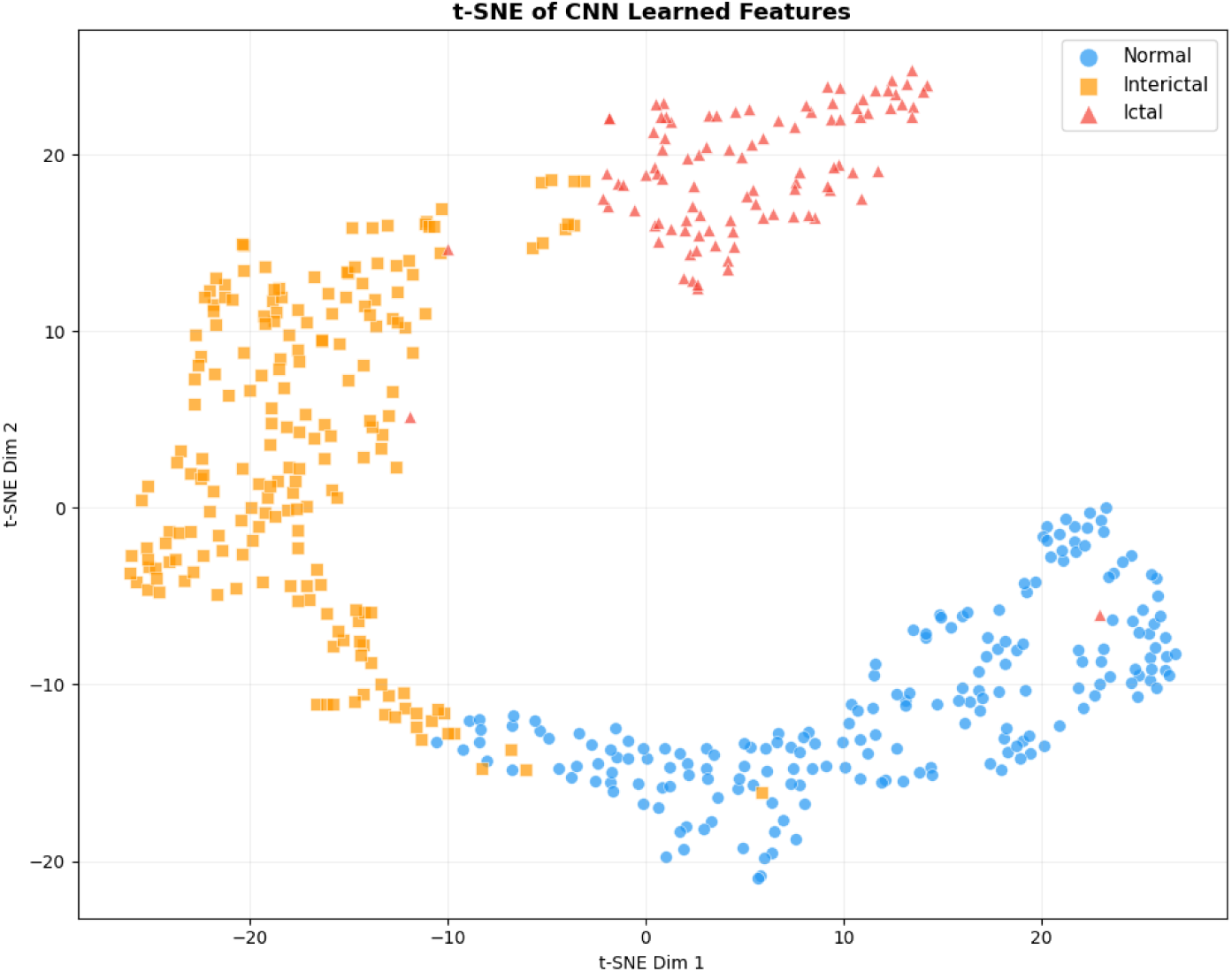
t-SNE of CNN features (baseline model, all 500 recordings).

**Figure 9.**
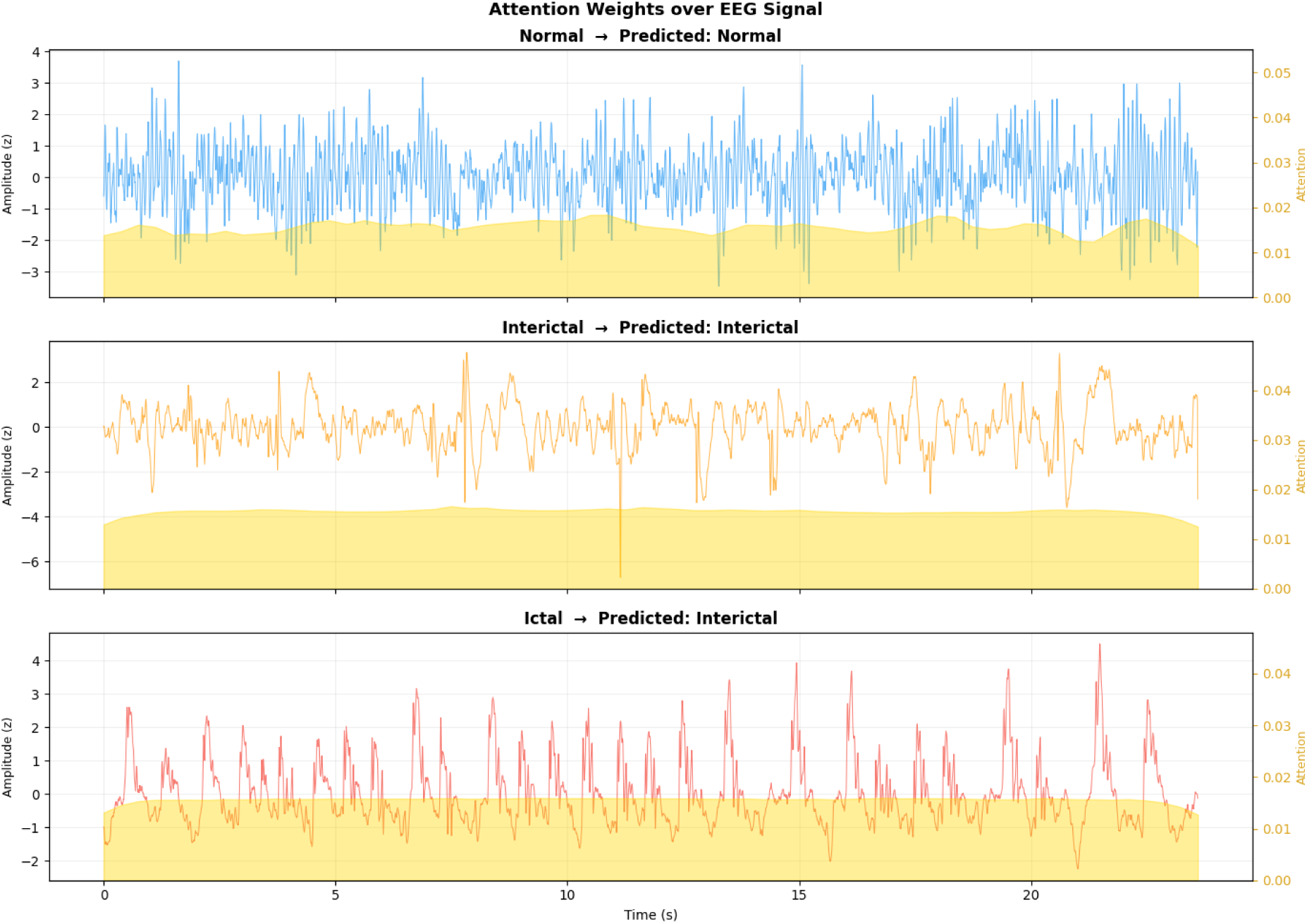
Attention weights (gold) over normalised EEG for the first test recording of each class. Predicted labels are in the panel titles; the Ictal example is misclassified as Interictal.

## 12 Discussion

DualBranchEEG combines raw-waveform and spectral views of EEG with a modest parameter budget. On the Bonn three-class task, both the CNN–BiLSTM baseline and the full model reach 96.00% held-out accuracy, and the dual-branch model attains the higher AUC-ROC (0.9952 vs. 0.9795). The class-wise band-power analysis confirms strong spectral separability, which makes the spectral branch plausible as a complementary input, but the ablation does not demonstrate that the extra branch or attention improves classification beyond what a CNN alone achieves on this data. Bonn is an easy, saturated benchmark; apparent differences of one recording cannot support architectural conclusions. The main practical value of this work is a transparent, reproducible reference implementation whose reported numbers match the executed code.

## 13 Future Work

- **Corrected pipeline.** Fit PSD standardisation on training data only, set Welch overlap explicitly, recompute the PSD from augmented signals, and rerun.
- **Rigorous comparison.** Budget-matched training, multiple seeds, nested cross-validation for every variant, and paired significance tests.
- **Larger datasets.** Multi-channel, patient-wise evaluation on CHB-MIT [24] and TUSZ [25].
- **Deployment.** Measure latency and memory on embedded hardware.
- **Interpretability.** Apply Grad-CAM to the final model on test data only and quantify saliency against annotated events.

## 14 Conclusion

We presented DualBranchEEG, a dual-branch network fusing a CNN–BiLSTM–attention temporal encoder with a Welch-PSD MLP for three-class EEG seizure classification. On the Bonn dataset it achieves 96.00% test accuracy, macro F1 0.9543 and macro AUC-ROC 0.9952, equalling the accuracy of a CNN–BiLSTM baseline (96.00%, AUC-ROC 0.9795; 98.40% in five-fold cross-validation). Ablations show no incremental accuracy gain from individual components, and the differences between variants are within sampling uncertainty. We document all deviations between the intended and executed protocol and identify the corrections needed before clinical or comparative claims can be made.

## Reproducibility Statement

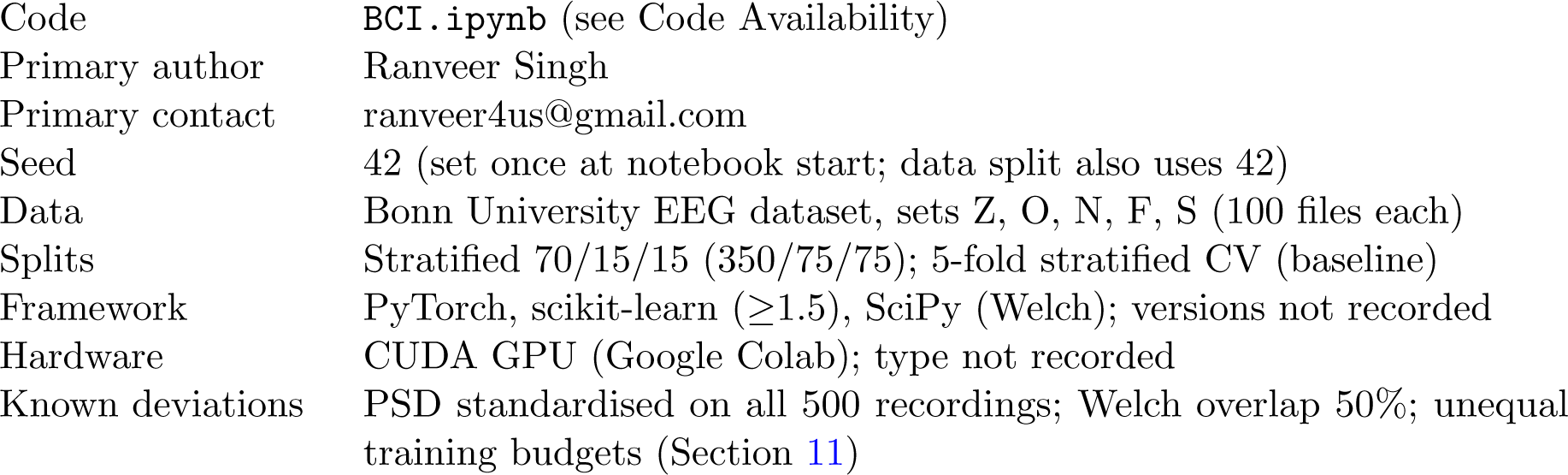

## Broader Impact

Automated seizure classification could reduce the burden of manual EEG review and support closed-loop neurostimulation. The present results come from a small, clean, single-channel benchmark and must not be interpreted as clinical performance. Deployment would require patient-wise validation on large multi-channel clinical datasets, prospective evaluation, calibration assessment and clinician oversight.

## Declarations

### Ethics statement

This study used only the publicly available Bonn University EEG dataset [12]. No new data were collected from human participants, and no additional ethical approval was required.

### Data availability

The Bonn University EEG dataset is publicly available from the Department of Epileptology, University of Bonn, and is described in [12]. **[Add the dataset download URL]**

### Code availability

The notebook used to produce all results (BCI.ipynb) is available at **[Add GitHub / Zenodo URL]**.

## A Supplementary Results and Hyperparameters

**Table 11.**
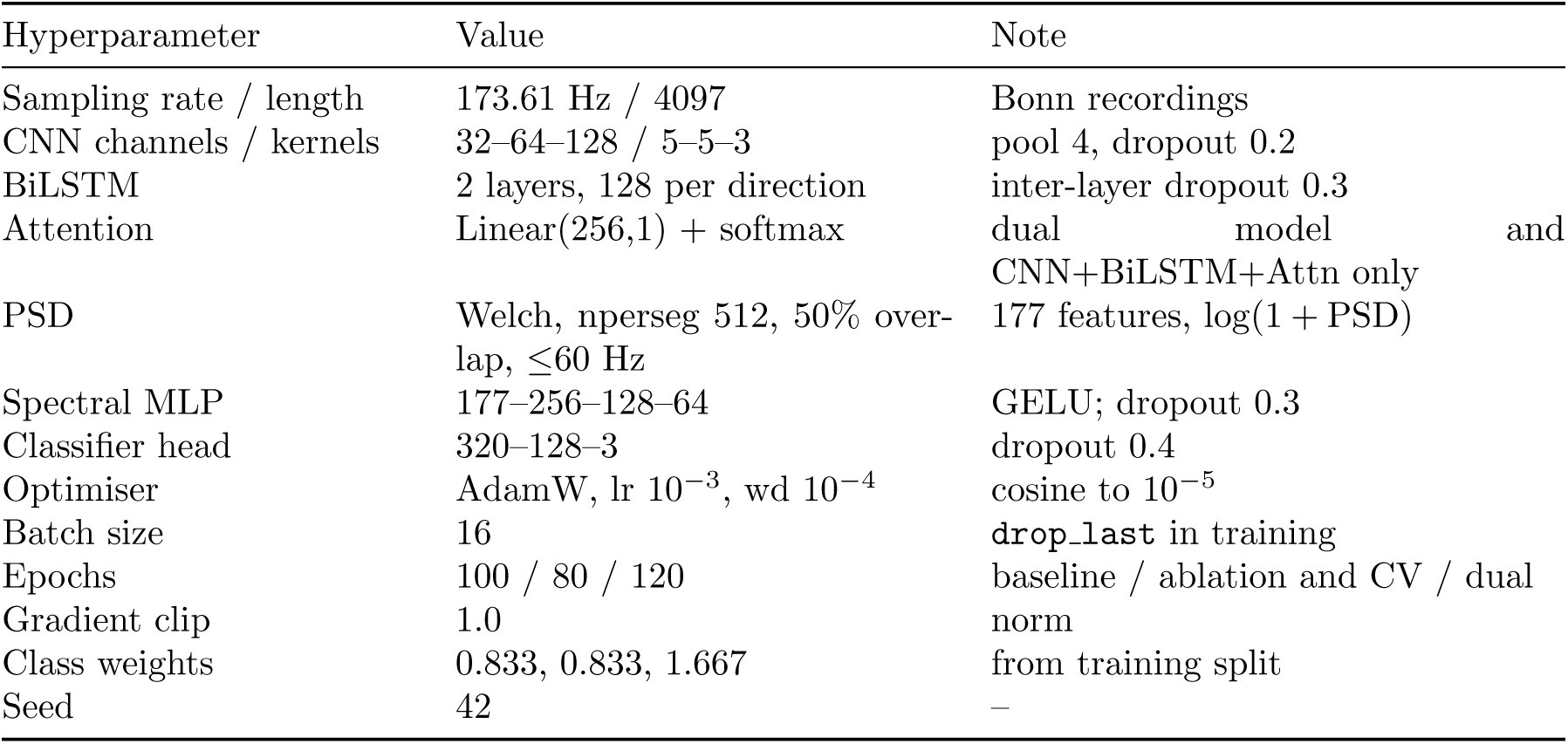
Hyperparameters as executed.

| Hyperparameter | Value | Note |
|---|---|---|
| Sampling rate / length | 173.61 Hz / 4097 | Bonn recordings |
| CNN channels / kernels | 32–64–128 / 5–5–3 | pool 4, dropout 0.2 |
| BiLSTM | 2 layers, 128 per direction | inter-layer dropout 0.3 |
| Attention | Linear(256,1) + softmax | dual model and CNN+BiLSTM+Attn only |
| PSD | Welch, nperseg 512, 50% over-lap, ≤60 Hz | 177 features, log(1 + PSD) |
| Spectral MLP | 177–256–128–64 | GELU; dropout 0.3 |
| Classifier head | 320–128–3 | dropout 0.4 |
| Optimiser | AdamW, lr 10<sup>−3</sup>, wd 10<sup>−4</sup> | cosine to 10<sup>−5</sup> |
| Batch size | 16 | drop_last in training |
| Epochs | 100 / 80 / 120 | baseline / ablation and CV / dual |
| Gradient clip | 1.0 | norm |
| Class weights | 0.833, 0.833, 1.667 | from training split |
| Seed | 42 | – |

**Figure 10.**
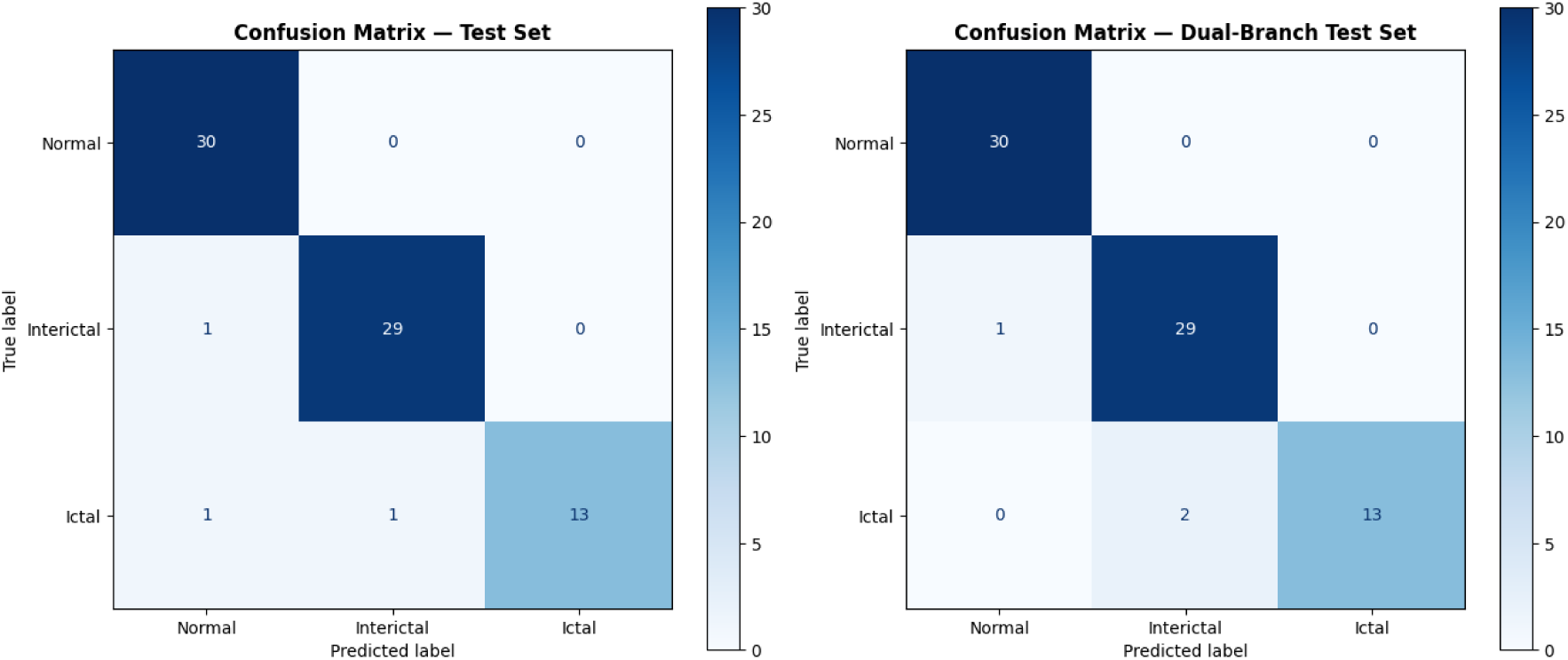
Test confusion matrices. Left: baseline CNN–BiLSTM; right: DualBranchEEG.

**Figure 11.**
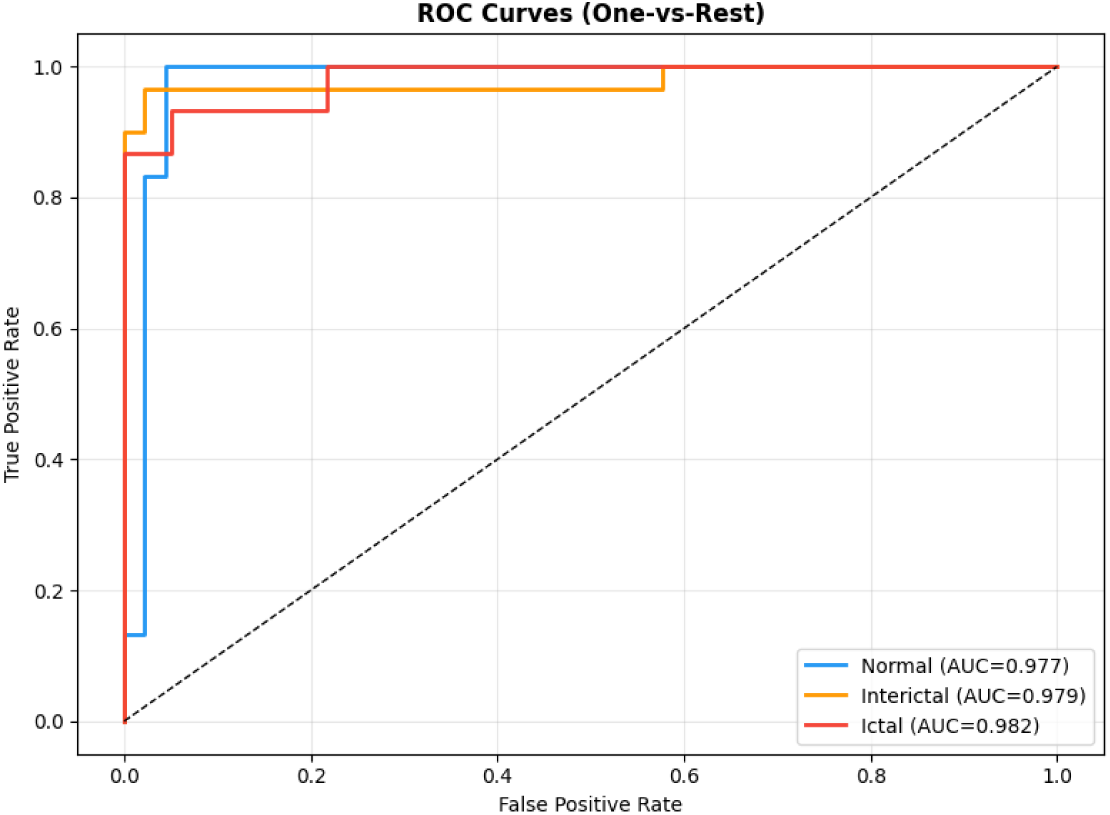
One-vs-rest ROC curves, baseline CNN–BiLSTM.

**Figure 12.**
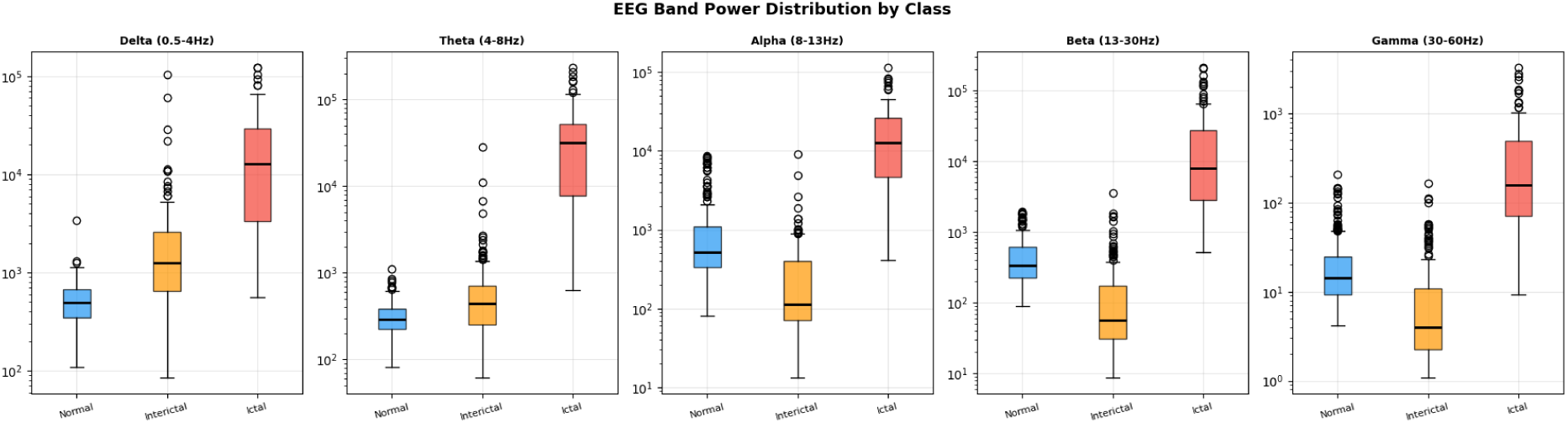
Band-power distributions by class (log scale). All pairwise differences in all five bands give *p <* 10*^−^*^5^ (Table 10).

**Figure 13.**
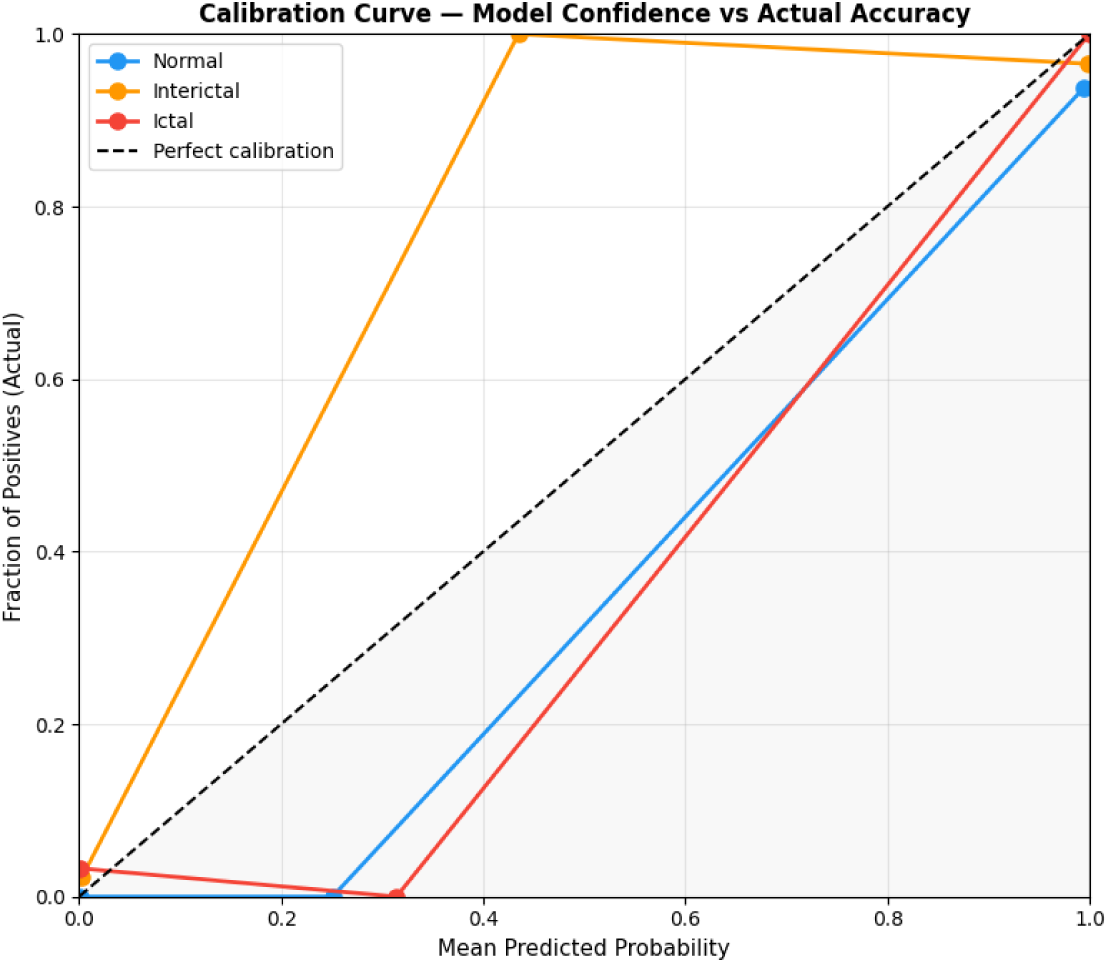
Calibration curves, baseline CNN–BiLSTM (one-vs-rest, 8 bins for the plotted curves; ECE values in Section 10.4 use 10 bins).

**Figure 14.**
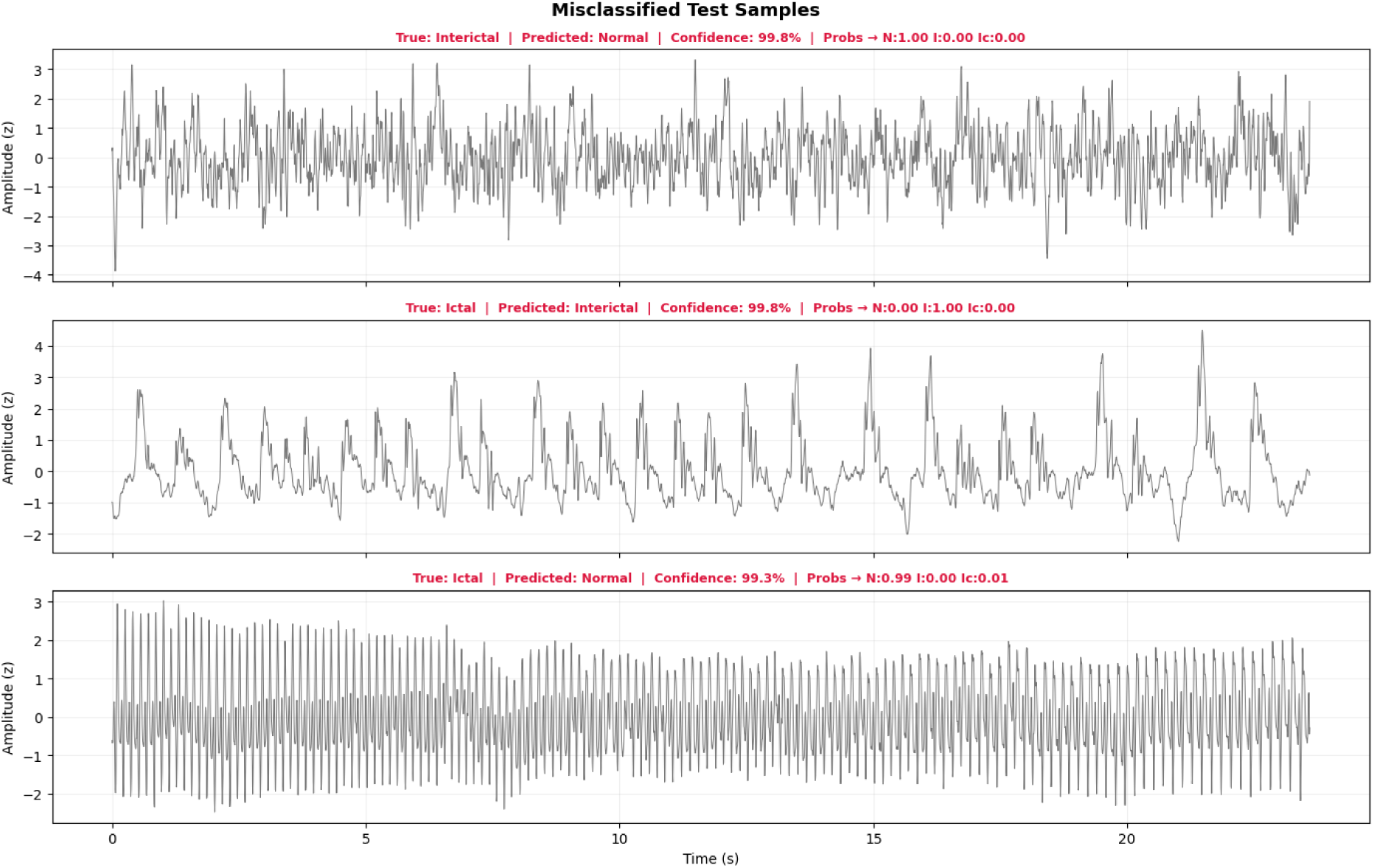
Misclassified test recordings, baseline CNN–BiLSTM.

